# Assessing the relationship between height growth and molecular genetic variation in Douglas-fir (*Pseudotsuga menziesii*) provenances

**DOI:** 10.1101/039818

**Authors:** Charalambos Neophytou, Anna-Maria Weisser, Daniel Landwehr, Muhidin Šeho, Ulrich Kohnle, Ingo Ensminger, Henning Wildhagen

## Abstract

Douglas-fir (*Pseudotsuga menziesii*) is a conifer tree native to western North America. In central Europe, it shows superior growth performance and is considered a suitable substitute for tree species impaired in vitality due to climate change. Maintenance and improvement of growth performance in a changing environment is a main challenge for forest tree breeders. In this context, genetic variation as a factor underlying phenotypic variation, but also as the basis for future adaptation, is of particular interest. The aims of this study were to analyse (i) genetic diversity of selected Douglas-fir provenances, (ii) variation in height growth among provenances, and (iii) to assess the link between genetic and phenotypic variation height growth. Genotyping was done on microsatellite loci. Effects of ‘provenance’, ‘genotype’, and ‘site’ on height growth were assessed by fitting mixed linear models. The most significant genetic differentiation was observed between provenances of the coastal variety, versus a provenance of the interior variety originating from British Columbia. Although genetic differentiation among provenances of the coastal variety was lower, genetic structures within this variety were identified. Moreover, genetic diversity showed a latitudinal gradient with the southernmost provenances being more diverse, probably reflecting the species' evolutionary history. The modelling approach revealed that height growth differed significantly by provenance, site, and the interaction between site and provenance, demonstrating that height growth is under strong genetic control. Additionally, this analysis showed that genetic variation captured by the genotyped microsatellite loci was significantly related to variation in height growth, providing statistical evidence for a genetic component in the observed phenotypic variation.

## Introduction

Douglas-fir (*Pseudotsuga menziesii* (Mirb.) Franco) is one of the most remarkable conifer species of western North American forests. It occurs naturally in a large geographic area ranging from northern latitude of 19° in Mexico to 55° in the Canadian province of British Columbia, being adapted to a wide variety of ecological conditions. Within its natural distribution area, it can attain majestic dimensions, reaching a height of more than 100 meters and a trunk diameter of more than 4.5 meters (Schütt et al. 2002, Aas 2008). Two varieties of Douglas-fir, the coastal (*P. menziesii* var. *menziesii*) and the interior (*P. menziesii* var. *glauca*) variety, can be well distinguished due to morphological, ecophysiological and growth differences. In addition, a series of transitional forms between these two varieties, as well as a large number of ecotypes can be found within the species’ range (Aas 2008, Konnert et al. 2008).

Due to its superior wood quality and high productivity, Douglas-fir has become one of the most important timber trees not only in North America, but also in several countries worldwide (e.g. throughout Europe, in New Zealand, Australia, Chile and Argentina), where it has been successfully introduced (Hermann and Lavender 1999). In these countries, its growth performance can exceed that of indigenous species. In Europe, for instance, Douglas-fir often displays higher volume increment and yield performance compared to economically significant forest tree species of local origin (Hermann and Lavander 1999), especially under adverse growth conditions (Eilmann and Rigling 2012). Therefore, it is of high interest as a source of timber in times of an increased demand on woody biomass.

Soon after the introduction of Douglas-fir to Europe, the factor ‘provenance’ was recognized as an important determinant for phenotypic traits (Kohnle et al. 2012, Šeho and Kohnle 2014).

Often, effects of ‘provenance’, i.e. seed source origin, on phenotypic traits are considered to be related to genetic variation among the respective provenances. Provenance research has revealed clinal variation in several adaptive traits like bud flushing, bud set, freezing tolerance and height growth along altitudinal and other climatic variables in Douglas-fir (Rehfeldt 1989, Leites et al. 2012, Chakraborty et al. 2015), but also in other conifer and broadleaved species (Aitken et al. 2008, Alberto et al. 2013, Kremer et al. 2014). However, only in recent years, this putative causal link was subject to systematic investigations in studies testing for an association between variation in phenotypes and allelic variation in candidate genes (for conifers e.g. González-Martínez et al. 2007, Eckert et al. 2009). The analysis of associations between phenotypic traits and functional genetic markers such as SNPs is complemented by the analysis of genetic diversity and stratification by means of putatively neutral genetic markers (i.e. occurring in non-expressed genomic regions), like microsatellite markers (Single Sequence Repeats; SSRs).

Microsatellites are appropriate markers for addressing a multitude of population genetic research questions. They are widely used in ecology and forest genetics (Selkoe and Toonen 2006, Guichoux et al. 2011, Martin et al. 2012). Due to their high variability, they provide insights into population genetic structures and evolutionary history of populations. On the other hand, genetic variation of SSR loci may also reflect selection and adaptation processes if these loci are located in the vicinity of genes subjected to natural selection. In this case, allele frequency changes at the gene region as a post-selection effect may also alter genetic differentiation at the SSR locus (Andolfatto 2001). Genome scans using a large number of loci have shown that divergent selection does not uniformly affect the genome, but is rather indicated by genome regions with an increased differentiation between populations, exceeding the neutral expectation. Specific microsatellite loci have often been classified as such ‘outlier’ loci, which is probably due to the indirect effects of natural selection (Scotti-Saintagne et al. 2004, Nosil et al. 2009, Tsumura et al. 2012). Moreover, recent SSR based genetic studies conducted in natural populations of forest tree species showed an association between genetic variation on the one hand and ecological (e.g. climatic or topographic parameters) or phenotypic variation (e.g. leaf size and growth) on the other (Ramírez-Valiente et al. 2010, Sork et al. 2010, Temunović et al. 2012).

In our study, we focused on a common garden experiment installed in Central Europe, in order to study the genetic and phenotypic variation among Douglas-fir provenances planted there. In particular, we aimed to analyse the effects of genotype and site conditions on growth performance of the provenances. Given that the source populations reside within a large geographic area, we expected to detect genetic differentiation due to isolation and drift among the provenances. Moreover, we hypothesize that genetic adaptation to variable local environments will be reflected in variation in height growth among the provenances. To test these hypotheses, we (i) analysed genetic diversity and differentiation among selected provenances based on data from SSR loci, (ii) we analysed the effects of the factor ‘provenance’ and of climate at provenance origins on variation in height growth, (iii) we assessed whether a provenance effect on height growth is related to genetic variation among the provenances and (iv) we discussed evolutionary implications by assessing statistical relationships between growth, molecular genetic variation, climate and geographic location of the provenances.

## Materials and methods

### Experimental plots and plant material

The sampled experimental sites are located in south-western Germany and were established in 1961 as part of an international provenance experiment with Douglas-fir (*Pseudotsuga menziesii* (Mirbel) Franco) with three-year-old saplings. Seeds had been collected from wind-pollinated parent trees in naturally regenerated stands in British Columbia (Canada), Washington, and Oregon (USA) (Strehlke 1959, Kenk and Thren 1984). From the various locations of the experimental series in south-western Germany, we selected along a gradient in elevation the experimental sites ‘Schluchsee’ (Dgl123; 1050 m above sea level (a.s.l.)), ‘Sindelfingen’ (Dgl91; 490 m a.s.l.), and ‘Wiesloch’ (Dgl122; 105 m a.s.l.). A brief description of these sites is presented in Table 1. We studied the coastal provenances ‘BC Cameron Lake’, ‘WA Darrington 3 Conrad Creek’, ‘OR Timber’ and ‘OR Santiam River’, as well as the interior provenance ‘BC Salmon Arm 31/102’. The selected provenances a wide latitudinal and longitudinal range of the North American seed source locations present at all three sites; for a detailed description of the origin of these provenances and site conditions, see Kohnle et al. (2012) and Kenk and Thren (1984). The latter reference includes a map showing the location of the experimental sites in Germany. Geographic positions of the provenances’ native habitats are presented in Figure 1. On each experimental site, each provenance was planted on two to four plots each of 0.1 ha size. Planting density was 3300 trees ha^−1^ at a spacing of 1.5 by 2 m. The arrangement of plots on each site did not follow either a systematic or random design in a strict sense, but plots were irregularly arranged so identical provenances never adjoined (Kohnle et al. 2012). A detailed description of the arrangement of the plots of the studied provenances is given in Supplementary Figure 1. The three selected sites are part of the network of long-term growth and yield experiments of the Forest Research Institute (FVA) Baden-Württemberg. The establishment and height-driven thinning regime of the plots is described in detail in Kohnle et al. (2012). The applied height-driven thinning regime resulted in rather uniform stand density dynamics. As a rule, the experiments displayed similar stand densities at identical stand heights. There were, however, some exceptions. As a consequence of the storm damage during the gales of 1990 and 1999, stand densities on some plots at the sites at ‘Wiesloch’ and ‘Sindelfingen’ were reduced below the height-specific target densities (see section *Collection of tree growth data* for details).

**Fig. 1.**
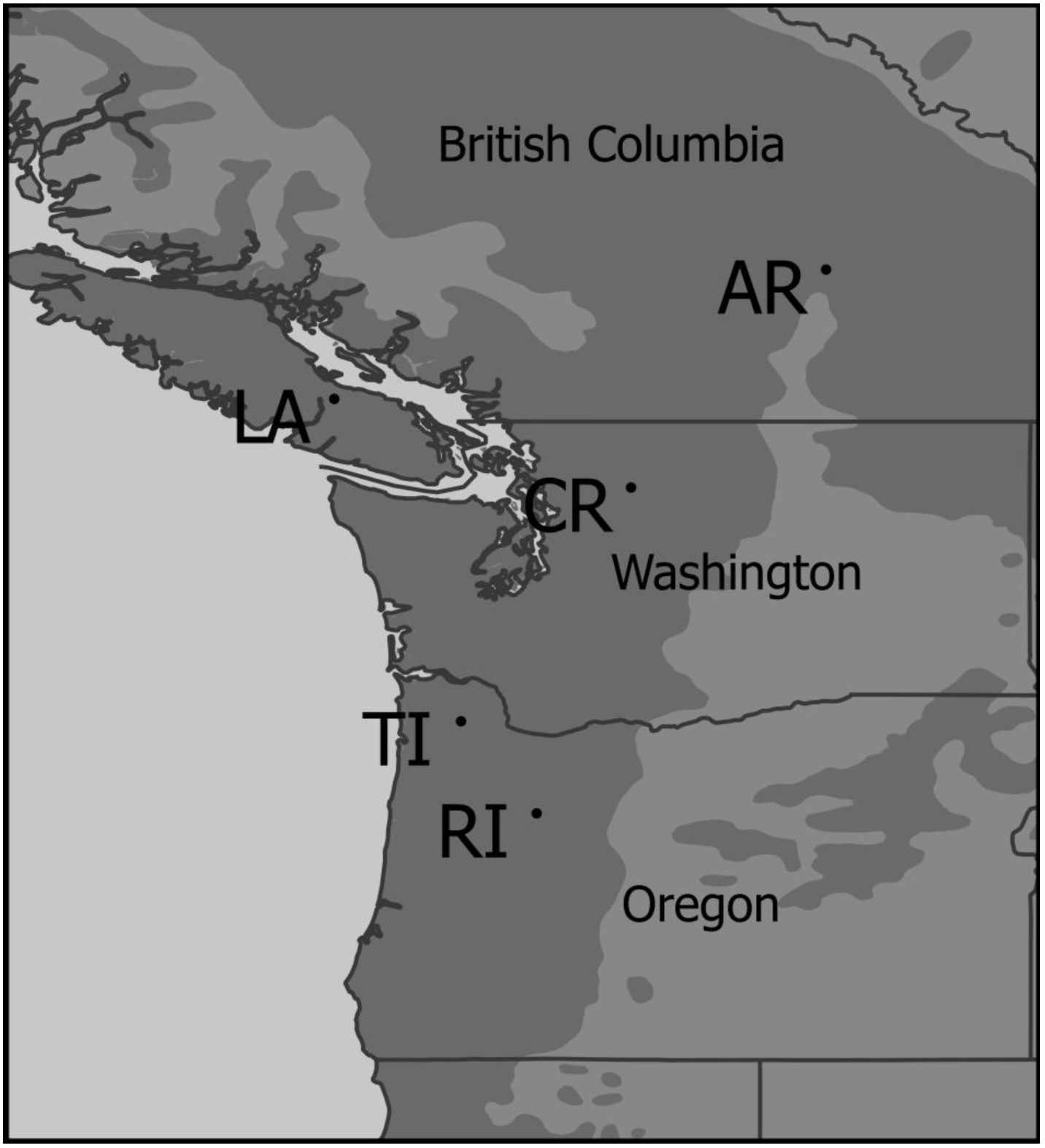
Map of the origin of provenances in British Columbia (Canada), Washington (USA) and Oregon (USA). Douglas-fir distribution range (Little 1971) is designated with dark grey (shapefile recovered by USGS publicly available at http://esp.cr.usgs.gov/data/atlas/little/). Shapefiles for political boundaries and coast lines were recovered by the website of the Commission for Environmental Cooperation of North America (CEC; data publicly available at http://www.cec.org/Page.asp?PageID=122&ContentID=2312&SiteNodeID=497&BL_ExpandID=). AR, Salmon Arm; CR, Conrad Creek; LA, Cameron Lake; TI, Timber; RI, Santiam River.

**Table 1.**
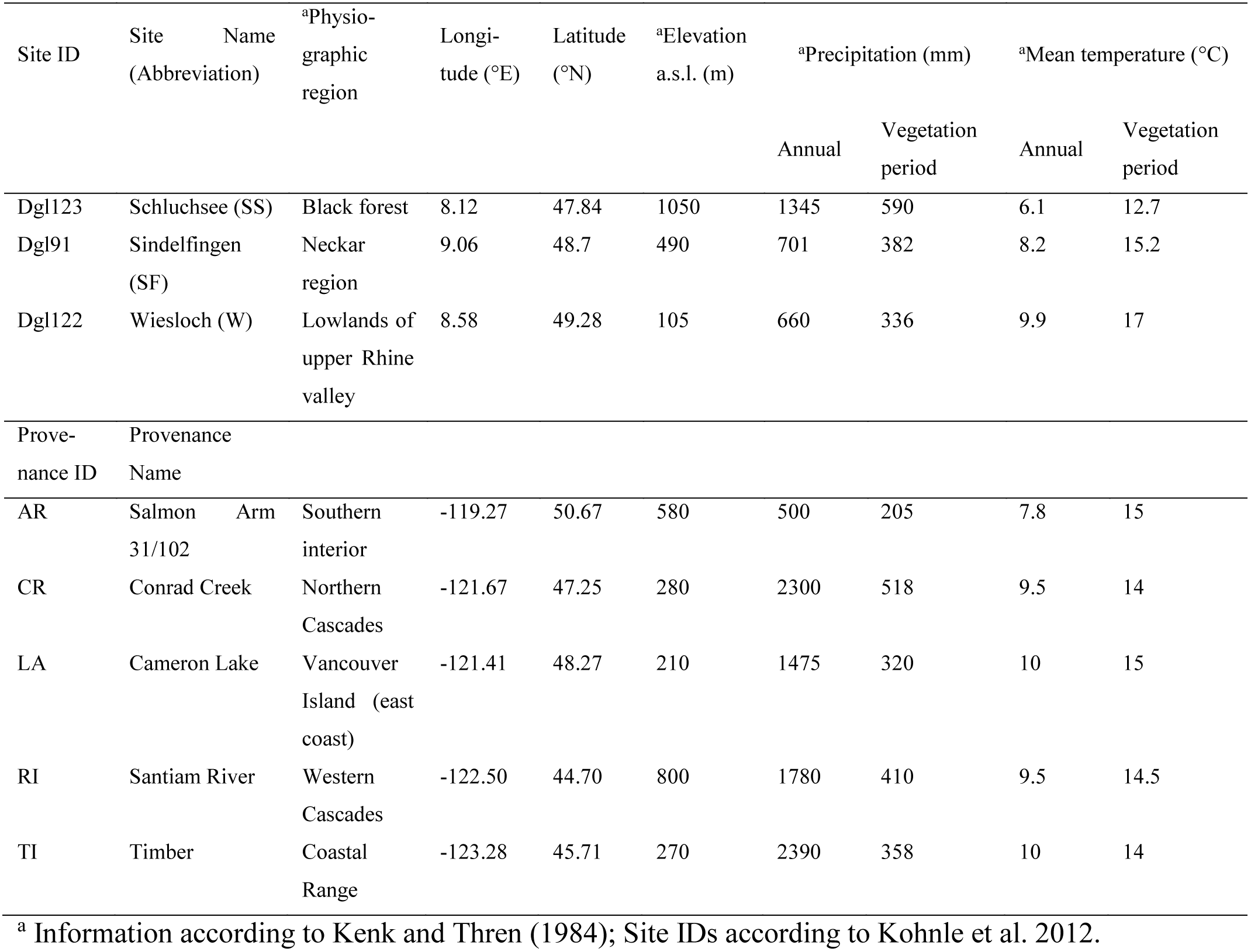
Characteristics of the three selected sites and of the source populations represented in these sites of the International Douglas-fir provenance trial

### Analysis of genetic diversity and differentiation

#### Sample collection for genotyping of microsatellite loci

For each provenance at each site needles or cambium was collected from 31 to 40 randomly chosen trees (see Supplementary Table 2 for the number of trees sampled per provenance and site).

These trees represent all plots per provenance and site, with the exception of provenance ‘Salmon Arm’ on site ‘Schluchsee’, where trees of only one of two plots were harvested. Until further utilisation the sampled material was stored at – 80°C.

#### Laboratory procedures

The needle and cambium tissue was first frozen in liquid nitrogen and subsequently homogenised in a mixer mill (MM30, Qiagen, Hilden, Germany). The DNA was extracted using the DNeasy^®^ 96 plant kit (Qiagen, Hilden, Germany) following the manufacturers’ protocol for extraction of frozen plant material with the following modifications: After grinding in lysis solution, samples were incubated for 40 min at 65 °C in a water bath; after the addition of the manufacturer’s precipitation buffer AP2, the samples were incubated for 30 min on ice instead of 10 min at −20 °C. The quantification of DNA was performed with a NanoDrop™1000 (Thermo Scientific, Karlsruhe, Germany). For genotyping, six unlinked and highly polymorphic dinucleotide simple sequence repeat (SSR) loci were amplified by polymerase chain reaction (PCR). The marker loci (PmOSU_1C3, PmOSU_2G12, PmOSU_3B2, PmOSU_3F1, PmOSU_3G9 and PmOSU_4A7) were selected from a previous study (Slavov et al. 2004). As in Krutovsky et al. (2009), we refer to these loci without the generic prefix “PmOSU”. PCRs were prepared using the components of the Multiplex PCR kit (Qiagen, Hilden, Germany) according to the manufacturers’ instructions. Oligonucleotide primers for the amplification of these loci were obtained from Applied Biosystems (Darmstadt, Germany) using the sequences given in Slavov et al. (2004) with the following set-up of fluorescent labelling: 6-Fam for 1C3, NED for 2G12, VIC for 3B2 and 4A7, PET for 3F1, NED for 3G9. The PCR program included an initial denaturation step at 95 °C for 15 min, 36 cycles at 94 °C for 30 s, an annealing step at 55 °C for 90 s, an elongation step at 72 °C for 45 s. Final elongation was performed at 60 °C for 30 min. The analysis of the fragment lengths was performed using the ABI-Prim^®^ 3100-Avant Genetic Analyser (Applied Biosystems, Darmstadt, Germany) by means of polymer 3100 Pop-4™, the Dye Set DS-33 and LIZ^®^ size standard (Applied Biosystems, Darmstadt, Germany).

#### Genotyping

The software GeneMapper^®^ version 4.0 (Applied Biosystems, Darmstadt, Germany) was used for genotyping. Allele bins were set by the software and manually corrected, if necessary. In order to check for rounding errors, the software TANDEM (Matschiner and Salzburger 2009) was used. This program hampers rounding errors that can occur during manual binning by applying a formula that rounds the allele sizes consistently (Matschiner and Salzburger 2009). Subsequently, data were checked for null alleles (non-amplifiable) and scoring errors (large allele drop-out and errors due to stuttering) using the software Microchecker v. 2.2.3 (Van Oosterhout et al. 2004) with the algorithm Brookfield 1. Within each locus, we applied 1,000 randomisations of alleles for the tests. The adjusted genotype lists were used for further analysis.

#### Statistical analysis of genetic diversity and differentiation among provenances

Observed (H_o_) and expected (H_e_) heterozygosity, as well as inbreeding coefficients (F_IS_; Weir and Cockerham 1984) were calculated using the software Genetix v. 4.05.2 (Belkhir et al. 2004). We defined 15 populations resulting from the combination of three sites and five provenances. Significance of F_IS_ within each population was tested by performing 1,000 random permutations of alleles over individuals using the same software. Number of alleles (n_a_), allelic richness (A) and pairwise fixation indices (F_ST_; Weir and Cockerham 1984) were computed using the software FSTAT v. 2.9.3.2 (Goudet 2001). Rarefaction size for the calculation of the allelic richness was set to 10. Following Goudet et al. (1996), a test of pairwise differentiation based on 1,000 random permutations of genotypes between populations was carried out with the same computer program. Given that multiple pairwise population comparisons were made, we applied a Bonferroni correction for multiple tests in order to find adjusted rejection probability values, following the instructions included in the software manual. The software Poptree2 (Takezaki et al. 2010) was used to calculate genetic distance D (Nei 1972) and for the assembly of a UPGMA phylogenetic tree. Reliability of clades was tested by bootstrapping with 1,000 repetitions.

In order to analyse genetic stratification and assign individuals to *K* subpopulations (genetic clusters), we employed a model-based Bayesian clustering analysis using the software STRUCTURE 2.3.3 (Pritchard et al. 2000). This software uses a Markov chain Monte Carlo procedure (MCMC) to infer unstructured subpopulations, which approach the Hardy-Weinberg equilibrium. Membership proportions of each individual to each one of *K* modelled subpopulations were calculated based on the individuals’ genotype. For the analysis, the admixture model was chosen which assumes that individuals may have mixed ancestry and could thus be assigned to more than one subpopulation. We used the locprior-function (Hubisz et al. 2009), which was developed to improve the clustering in situations where individuals are sampled from different geographic locations. We assigned the 15 populations to the five provenances. 100,000 updates of the Markov chain and 100,000 MCMC iterations were performed. The procedure was run for *K* = 1, …, 15. Ten runs were performed for each *K*. To detect the uppermost hierarchical level of structure and to provide an accurate estimation of population clustering the parameter Δ*K* after Evanno et al. (2005) was calculated with the software Structure-Harvester (Earl and vonHoldt 2012). The maximal Δ*K* was found for two assumed subpopulations (*K* = 2). The second highest Δ*K* was found for of *K* = 3.

To find the optimal alignment of the replicate cluster analysis (10 runs for each *K*) the software CLUMPP v. 1.1.2 (Jakobsson and Rosenberg 2007) was applied. The output from CLUMPP was used as input for the cluster visualisation program DISTRUCT v.1.1 (Rosenberg 2004).

To validate our results from clustering with STRUCTURE, we used an alternative way to detect structures by applying a multivariate statistical method suited for the detection of clusters and for the reduction of multivariate data to a smaller number of dimensions. For this purpose, single locus genotypes are transformed into allelic variables by assigning scores of 0, 1, or 2 for individuals not carrying a particular allele, carrying it in a heterozygous state, or a homozygous state, respectively (as proposed by She et al. 1987 and Duplantier et al. 1990). A widely used method for the identification of population stratification, which accommodates the discrete nature of the allelic variables is correspondence analysis. This analysis is implemented in the software Genetix v. 4.05.2 (Belkhir et al. 2004), which is frequently used in population genetics (e.g. Vasemägi et al. 2001, Belaj et al. 2007, Bryja et al. 2010). We opted to apply such a factorial correspondence analysis (FCA) to our data by employing the function “AFC sur populations” of the Genetix software, where the predefined populations represented by their allele frequency vectors are used as the object for the derivation of the new composite variables (factors) and individuals are subsequently given new coordinates within this composite hyperspace.

Besides their utility for identification of genetic structures, composite variables derived by multivariate analyses may also better reflect natural selection and adaptation processes than allele frequencies themselves (Grivet et al. 2008, Sork et al. 2010). This is partly because adaptation processes often involve polygenic traits causing moderate effects across the genome (Pritchard et al. 2010).

### Analysis of tree height data in relation to ‘site’, ‘provenance’, ‘genotype’, and climate

#### Collection of tree growth data

The long-term experiments have been routinely monitored by periodic re-measurements in intervals of usually five years commencing at a stand height of approximately 10 m (stand age 18-22 years). The last measurement took place in winter 2011/2012 when the stands were 54 years old; data presented here originate from this last measurement. The height of the trees was determined as the average of two hypsometer measurements (Vertex IV, Haglöf, Langsele, Sweden). In addition to the trees measured in winter 2011/2012, height data of trees cut for a study on branch and stem characteristics, measured in winter 2010/2011 were included in the analysis. The height of these trees was adjusted by adding the annual height increment measured for the last growing season to the determined height. The increment of the last growing season was defined as the length from the tip of the crown to the topmost whorl. All further data analyses on possible provenance and site specific influences on height growth were exclusively based on these measured or projected height data. In total, among the 598 trees measured individually for height, the height of 63 trees was estimated through projection of the height increment of the last year (see Supplementary Table 1 for a summary of the number of trees measured for height per provenance and site).

As it is the rule in even-aged stands, the social (competition) status affects the expression of tree height in the stands of the trial: thick, dominant trees tend to be taller than co-dominant and suppressed trees with smaller diameter. This correlation is captured by the stand height curve. As we were using height data from trees of different diameter, we had to account for possible confounding diameter-related impacts on height. For this purpose, we derived plot-specific height curves by (i) measuring all trees present on the plot for diameter at breast height (at about 1.3 m height; dbh) by averaging two rectangular calliper measurements to the closest mm, and (ii) by measuring approximately 30 trees selected across the range of the dbh for height. From these measurements, the plot specific height curves were derived using the program and function described by Ehring et al. (1999).

Since the diameters of all trees were measured, stand-specific diameter characteristic could be calculated from the individual diameters: for example, d200 is calculated as the diameter of the mean basal area tree of the 200 thickest trees ha^−1^ (ca. 20 trees per plot). Corresponding stand-specific height characteristics can then be calculated from the plot-specific height curve (e.g. h200 as the height curve value corresponding to d200). Based on these plot-specific height growth curves, we were able to account for possible competition-dependent effects on tree heights: firstly, we restricted all further analyses to height measurements of the dominant trees of the d200 collective (200 thickest trees·ha^−1^; restriction to dominant trees was based on the rationale, that the trajectory of the stand height curve is usually relatively flat among the dominant trees of a stand). Secondly, we developed a correction term (cor_h_*status*_) that was added to a tree’s actually measured height. Based on the stand height curves constructed from the measurements in winter 2011/2012, this correction term calculates as follows:

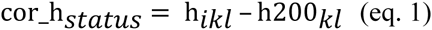

where *h*_*ikl*_ is the height of tree *i* on plot *k* at site *l* derived from the plot-specific stand height curve and the *dbh* of tree *i* and h200_*kl*_ is the height of the d200 tree on plot *k* at site *l* derived from the plot-specific stand height curve and d200.

A number of plots on the sites ‘Wiesloch’ and ‘Sindelfingen’ had suffered from wind-throw damage during gales in 1990 or 1999, respectively. In tendency tall trees had suffered more as to be expected from general models on storm damage demonstrating an increase of damage probability with increasing tree height (Schmidt et al. 2010, Albrecht et al. 2012, 2013): based on the height data obtained from the measurement prior to the storm, h200 of the trees present before the storm and h200 of the trees actually surviving the storm revealed a reduction of stand height ranging between 0 and 1.4 m with an average of 0.41 m and a standard deviation of 0.43 m. On plots where such storm damage-induced reductions of h200 exceeded 0.1 m (on 22 out of 33 plots), we added another correction term (cor_h_*storm*_) as a plot-specific offset value to the individual tree height data in order to compensate for the damage-induced effects on stand height:

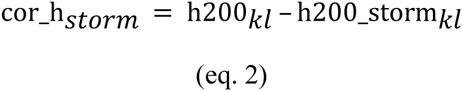

where h200_*kl*_ is the h200 of plot *k* at site *l* calculated from the plot-specific height curve based on all trees present at the measurement before the storm, and h200_storm_*kl*_ is h200 of plot *k* at site *l* calculated from the plot-specific height curve based on the trees actually surviving the storm until the last measurement. For the 22 plots where a correction term was introduced cor_h_*storm*_ ranged from 0.1 – 1.4 m and averaged 0.61 m (standard deviation: 0.38 m). Statistical analyses of the proportion of the cumulated damage-related removals of the plots’ total basal area production and of the correction term cor_h_*storm*_ revealed that the damaging effect was not significantly different among provenances (data not shown).

#### Statistical analysis of tree height data in relation to ‘site’, ‘provenance’, and ‘genotype’

Since not all genotyped trees were scored for height growth, we first matched the genotype and height growth data sets, yielding 315 complete observations (see Supplementary Table 3 for the number of matching trees per provenance and site). Normal distribution of this set of height data was verified graphically. Linear mixed-effects models estimating the height of individual trees were fitted with the function ‘lme’ provided within the ‘nlme’ library (Pinheiro et al. 2012) within unix version 2.14.0 of the statistical software environment R (R Development Core Team 2011). The aim of the modelling approach was two-fold: First, we wanted to assess if height growth differed among provenances and if these differences depend on the factor ‘site’. Second we wanted to test if differences in height growth found among populations are related to genetic variation in SSR loci. To pursue these two aims the modelling was done in two independent approaches: To address the first aim, the factors ‘provenance’, ‘site’, as well as the interaction between ‘provenance’ and ‘site’ were included as fixed explanatory factors in the model (referred to as the provenance-based model). To address the second aim, another modelling procedure was done where we omitted ‘provenance’ but included the membership proportions of individual trees in *K* = 5 clusters as calculated by STRUCTURE (see above for details of this analysis), and the factor ‘site’ (referred to as the STRUCTURE-based model).

Following Zuur et al. (2007), we started the model building process with full, potentially beyond-optimal models, including a random intercept for ‘plot’ as a random effect. Next, appropriate fixed-effects structures were selected by subsequently dropping non-significant fixed effects (P > 0.05) with highest P-values as computed with the function ‘anova’. Parameter estimation was computed with the maximum-likelihood technique. The reduced model was compared to the respective full model using BIC. This procedure was iterated until all factors retained in the model were significant. The parameters for the final models were re-estimated with the restricted-maximum-likelihood method.

In the provenance-based approach including ‘provenance’ and ‘site’, the height of tree *i* belonging to provenance *j* growing on plot *k* at site *l* was modelled as

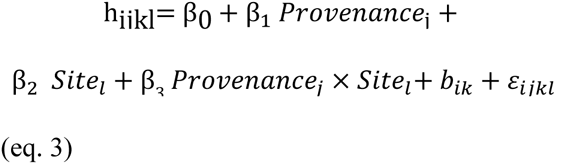

with *i* = 1, …, 315; *j* = 1, …, 5; *k* = 1, …, 31; *l* = 1, 2, 3. The fixed effects are represented by β_0_ (intercept), β_1_ (slope for ‘provenance’), β_2_ (slope for ‘site’) and β_3_ (slope for interaction ‘provenance × site’).

In the second modelling approach omitting ‘provenance’ but including memberships to the genetic clusters identified by STRUCTURE under *K* = 5 (STRUCTURE-based model), model selection was started with the following full model describing the height of tree *i* growing on plot *k* at site *l* as:

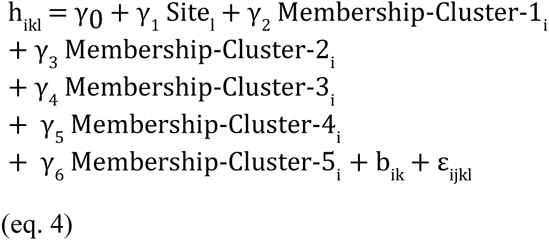

with *i* = 1, …, 315; *k* = 1, …, 31. The fixed effects are represented by γ_0_ (intercept), γ_1_ (slope for membership in cluster 1), γ_2_ (slope for membership in cluster 2), γ_3_ (slope for membership in cluster 3), γ_4_ (slope for membership in cluster 4), γ_5_ (slope for membership in cluster 5) and γ_6_ (slope for ‘site’).

Model selection resulted in a model describing the height of tree *i* growing on plot *k* at site *l* as:

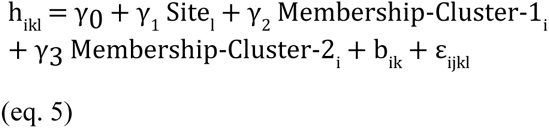

with *i* = 1, …, 315; *k* = 1, …, 31. The fixed effects are represented by γ_0_ (intercept), γ_1_ (slope for membership in cluster 1), γ_2_ (slope for membership in cluster 2) and γ_3_ (slope for ‘site’).

For both models, the random effect for ‘plot’ is denoted by b_ik_. Within-group errors are assumed to be independent identically normally distributed.

Random effects are assumed to be normally distributed. Within-group errors and random effects are assumed to be independent.

Correlation between observations of the same level of ‘plot’ was calculated as

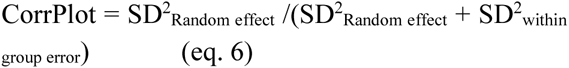

where SD denotes the standard deviation.

The validity of the final models was evaluated by graphically inspecting the residuals for normality, constant variance and centring around zero (Supplementary Figure 3, 4). The assumption of normally distributed random effects was assessed graphically, as well (Supplementary Figure 5). The explanatory power of the models was calculated as the squared correlation between the observed values and the predicted values based on fixed-effects and random effects.

In order to compare height growth among provenances within each site, separate models for each site including only ‘provenance’ as a main effect were fitted.

In addition, we applied Mantel tests and partial Mantel tests as a further approach to study the relationship between growth, on the one hand, and site condition or genetic differentiation on the other. For this purpose, we computed dissimilarity matrices comparing all 15 populations pairwise for all these three factors. For calculating growth (phenotypic) dissimilarity, we used height and dbh. After standardising both variables (using the function ‘scale’ in R), we calculated Euclidean distances between populations using the function ‘dist’ in R. In the same way, we produced a matrix of climatic dissimilarity among sites based on all four climatic variables presented in Table 1. As a matrix of genetic dissimilarity, we used Nei distances calculated with the software POPTREE2. We performed the Mantel Tests using the functions ‘mantel’ and ‘partial.mantel’ of the package vegan 2.0-10 (Oksanen et al. 2013) based on 9999 permutations and Pearson’s correlation coefficient.

#### Statistical analysis of tree height data in relation to climatic conditions at the provenances origin

Finally, we tested whether there are effects of evolutionary processes from the place of origin on growth performance or genetic variation. Following the procedure described above, we studied the correlation between the matrices of (1) genetic differentiation or (2) phenotypic on the one hand and climatic or geographic differentiation among provenances on the other. In the first case, we aimed to analyse whether there is an effect of isolation by distance (IBD) or isolation by adaptation (IBA) on genetic variation at putatively neutral loci. In the second case, we aimed to test whether there are similar effects on growth, a trait of adaptive relevance.

## Results

### Genetic diversity across loci

Two of the six loci could not be used for the population genetic analysis. Although we attempted to optimise PCR conditions for the primers of locus 1C3, no evaluable electropherogram could be generated. Besides excessive stuttering, additional peaks, corresponding to additional PCR amplicons, occurred in a large number of samples at this locus. For locus 3B2 no bin-set could be generated. Fragment sizes displayed a rather continuous distribution, whereas a systematic shift of two base pairs would be expected for this locus. Thus, an assignment of the observed peaks to different allele clusters (bins) was not possible and the locus was removed from further analyses. For the remaining loci, genotype lists were adjusted using the software Microchecker 2.2.3, in case of scoring errors (Van Oosterhout et al. 2004). Genotyping errors detected by Microchecker included ‘null’ alleles and excessive stuttering (Supplementary Table 4).

An analysis of the variability of the four analysed loci over all populations after genotypic adjustments for null alleles Revealed that the number of alleles per locus (without considering null alleles) ranged from 32 at locus 3G9 to 50 at locus 4A7 (Table 2). For the loci 3F1, 2G12 and 4A7 allelic richness was equal or higher than 12.04, while at locus 3G9 it was only 9.85 (Table 2). The observed heterozygosity (H_o_) varied from 0.724 to 0.857. Expected heterozygosity (H_e_) was generally high (at least 0.887). Even after genotypic adjustments for null alleles, inbreeding coefficient F_IS_ averaged over all loci was 0.145.

**Table 2.** Diversity measures for each locus in the whole sample after genotype adjustments for null alleles. Genotype adjustments for null alleles were performed with Microchecker v. 2.2.3 (van Oosterhout et al. 2004). Number of alleles per locus (n_a_) and allelic richness (A; rarefaction size = 10) were calculated with FSTAT v. 2.9.3.2 (Goudet 2001), observed (H_o_) and expected heterozygosity (H_e_), inbreeding coefficient (F_IS_) with Genetix v. 4.05.2 (Belkhir et al. 2004) and null allele frequencies (f_0_) with Microchecker v. 2.2.3

| Locus | $n_a$ | A | $H_o$ | $H_e$ | $F_{IS}$ | $f_0^a$ |
| --- | --- | --- | --- | --- | --- | --- |
| 3F1 | 39 | 12.51 | 0.784 | 0.940 | 0.167 | 0.120 |
| 2G12 | 41 | 12.04 | 0.857 | 0.933 | 0.082 | 0.042 |
| 4A7 | 50 | 12.77 | 0.724 | 0.930 | 0.223 | 0.313 |
| 3G9 | 32 | 9.85 | 0.752 | 0.887 | 0.152 | 0.258 |
| Average | 40.5±3.7 | 11.7±0.6 | 0.78±0.03 | 0.92±0.01 | 0.16±0.03 | 0.18±0.06 |

<sup>a</sup> = Brookfield 1 estimate was used for null allele frequencies

The lowest F_IS_ value was found for locus 2G12 and the highest value for 4A7 (Table 2). Given the high estimated frequencies of null alleles, only adjusted genotypic data were used for further genetic analyses.

The genetic diversity of the five study provenances is presented in Table 3. With the exception of H_o_, the highest diversity values were measured for provenances ‘Santiam River’ and ‘Timber’. In general, these two provenances showed relatively high diversity across loci. In the remaining three provenances, high diversity values were observed only at specific loci. For instance, provenance ‘Conrad Creek’ displayed high diversity at locus 3G9 and provenance ‘Salmon Arm’ at locus 2G12 (data not shown). On average, the provenances ‘Salmon Arm’ (AR) and ‘Cameron Lake’ (LA) showed the lowest diversity values. However, H_o_ for ‘Cameron Lake’ was the highest due to low heterozygote deficiency marked by the lowest inbreeding coefficient (F_IS_).

**Table 3.** Diversity measures for each provenance averaged over loci and field sites. Average number of alleles per locus (n_a_) and allelic richness (A; rarefaction size = 10) were calculated with FSTAT v. 2.9.3.2 (Goudet 2001), while observed (H_o_) and expected heterozygosity (H_e_), inbreeding coefficient (F_IS_) were calculated with Genetix v. 4.05.2 (Belkhir et al. 2004). Abbreviations for provenances: AR, ‘Salmon Arm’; CR, ‘Conrad Creek’; LA, ‘Cameron Lake’; RI, ‘Santiam River’; TI, ‘Timber’. Abbreviations for sites: W, ‘Wiesloch’, SS; ‘Schluchsee’; SF, Sindelfingen.

| Site | N | $n_a$ | A | $H_o$ | $H_e$ | $F_{IS}$ |
| --- | --- | --- | --- | --- | --- | --- |
| AR | 110 | 23.25 | 10.55 | 0.758 | 0.892 | 0.155 |
| CR | 118 | 23.75 | 11.06 | 0.761 | 0.909 | 0.167 |
| LA | 111 | 22.50 | 10.23 | 0.811 | 0.897 | 0.101 |
| RI | 113 | 29.25 | 11.79 | 0.775 | 0.920 | 0.162 |
| TI | 108 | 28.50 | 11.74 | 0.788 | 0.923 | 0.150 |
| Mean |  | 25.45±1.42 | 11.07±0.31 | 0.784±0.010 | 0.912±0.006 | 0.147±0.012 |

### Genetic variation, stratification, and differentiation among populations

F_ST_ values between pairs of provenances are presented in Table 4. The most significant differentiation in terms of pairwise F_ST_ values was found between the provenance ‘Salmon Arm’ (AR) on the one hand and the other four provenances on the other (Table 4).

**Table 4.** Pairwise F_ST_-values among provenances (below diagonal) and probability based on test of differentiation (above diagonal) according to Goudet et al. (1996) computed with FSTAT v. 2.9.3.2 (Goudet et al. 2001). The indicative adjusted nominal level (5%) for multiple comparisons after Bonferroni correction was 0.005. Abbreviations for provenances (Prov.): AR, ‘Salmon Arm’; CR, ‘Conrad Creek’; LA, ‘Cameron Lake’; RI, ‘Santiam River’; TI, ‘Timber’. Abbreviations for sites: W, ‘Wiesloch’, SS; ‘Schluchsee’; SF, Sindelfingen.

| Provenance | AR | CR | LA | RI | TI |
| --- | --- | --- | --- | --- | --- |
| AR |  | 0.000 | 0.000 | 0.000 | 0.000 |
| CR | 0.023 |  | 0.000 | 0.000 | 0.000 |
| LA | 0.029 | 0.008 |  | 0.000 | 0.000 |
| RI | 0.022 | 0.007 | 0.009 |  | 0.164 |
| TI | 0.025 | 0.007 | 0.011 | 0.000 |  |

Genetic variation among the 15 populations was analysed by computing genetic distances after Nei (1972). Genetic distances between the populations were low when pairs of the same provenance (but from different sites) were considered (highest D = 0.37, Supplementary Table 5). For most pairs involving one of the populations of provenance ‘Salmon Arm’, the genetic distances were high (lowest D = 0.29, highest D = 0.69, Supplementary Table 5). To visualise and statistically validate the genetic distances between the populations a phylogenetic tree was constructed (Supplementary Figure 1). It shows that the main division (94 % of the bootstraps) is the branch of the three populations of the provenance ‘Salmon Arm’ (AR), whereas all populations of the other four provenances are clustered on a second main branch.

Figure 2 shows the genetic structure of the populations under the assumption of *K* = 2, …, 5 as computed with the software STRUCTURE (Pritchard et al. 2000). For *K* = 2 obviously the individuals of the provenance ‘Salmon Arm’ at all field sites are assigned to one subpopulation whereas the individuals of the four other provenances over all field sites were assigned together to the second subpopulation. In both cases, membership coefficients were high. The genetic structure assuming *K* = 3 shows once again that trees of the provenance ‘Salmon Arm’ are clearly assigned together to one subpopulation (marked blue) with high membership coefficients. Individuals of the provenance ‘Cameron Lake’ were assigned to a second subpopulation with little admixture with other clusters. A third subpopulation was mainly represented by the provenances ‘Santiam River’ and ‘Timber’, which showed a limited degree of admixture with the second subpopulation. Trees of the provenance ‘Conrad Creek’, displayed mixed coancestry, with equal membership to both aforementioned clusters. By setting *K* = 4, provenances ‘Santiam River’ and ‘Timber’ constitute a fourth cluster with comparably low admixture. No further structure was revealed by setting *K* = 5 or higher.

**Fig. 2.**
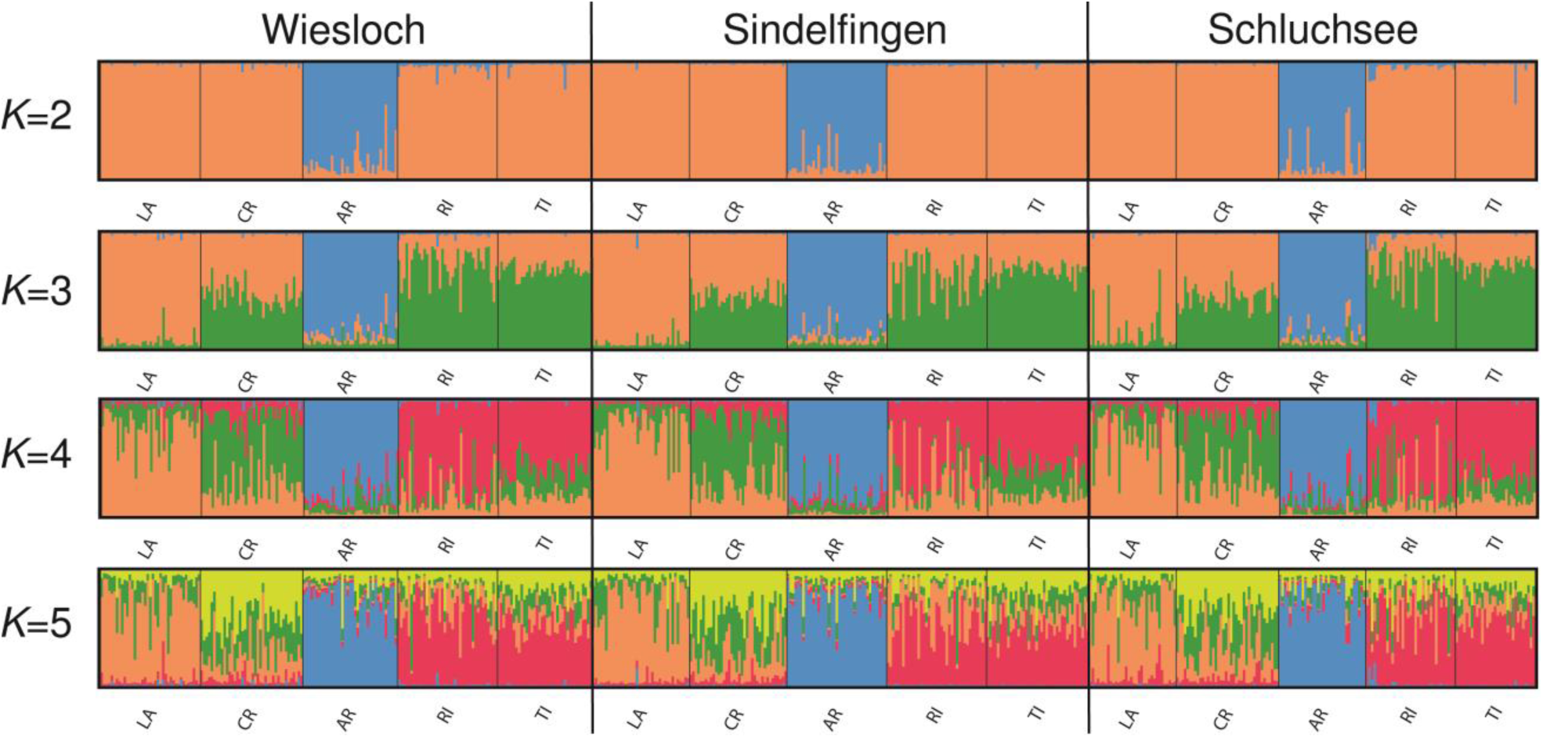
Genetic structure Douglas-fir trees of five provenances growing on three site in south-western Germany computed on the basis of four microsatellite loci for *K* = 2 up to *K* = 5 assumed subpopulations. With the software STRUCTURE 2.3.3 (Pritchard et al. 2000) the 15 populations and their individuals were assigned to the subpopulations. Each individual is represented with a vertical bar and each inferred cluster is marked with a different colour (colours are visible only in the online version of this article). Abbreviations for provenances: AR, ‘Salmon Arm’; CR, ‘Conrad Creek’; LA, ‘Cameron Lake’; RI, ‘Santiam River’; TI, ‘Timber’.

To validate the results of population structure calculated by STRUCTURE, we analysed the genotype data with a non-model-based factorial correspondence analysis (FCA) as implemented in the software Genetix (v. 4.05.2, Belkhir et al. 2004). The first four factors accounted for 23.17, 12.17, 9.02 and 8.48% (a cumulative 52.84%) of the total variation respectively. The most prominent pattern detected with this analysis was a separation along the first axis (Factor-1, see Supplementary Figure 6): Trees of the provenance ‘Salmon Arm’ showed higher scores on this axis, while individuals of the other four provenances clustered around a score of zero. There was a tendency for a separation of provenances ‘Santiam River’ and ‘Cameron Lake’ along Factor-2 (Supplementary Figure 6). Trees of both provenances, ‘Santiam River’ and ‘Salmon Arm’, were spread over a wider range of Factor-3 compared to individuals of the other provenances (Supplementary Figure 6).

### Height of Douglas-fir trees as affected by ‘provenance’, ‘site’ and ‘genotype’

Exploratory analysis of data on height growth of Douglas-fir trees approximately at the age of 50 years indicates effects of site, provenance, and interaction between these two factors (Figure 3). Averaged over the three sites investigated in this study, trees of provenance ‘Cameron Lake’ reached a height of 30.7 ± 2.2 m (mean ± standard deviation), those of provenance ‘Conrad Creek’ 32.2 ± 2.3 m, of provenance ‘Salmon Arm’ 30.1 ± 2.2 m, of provenance ‘Santiam River’ 30.4 ± 2.3 m, and those of provenance ‘Timber’ 31.7 ± 1.8 m. The mean heights of trees on the three sites averaged over provenances were 32.7 ± 1.4 m for site ‘Wiesloch’, 29.0 ± 1.8 m for site ‘Schluchsee’, and 30.6 ± 2.2 m for site ‘Sindelfingen’.

**Fig. 3.**
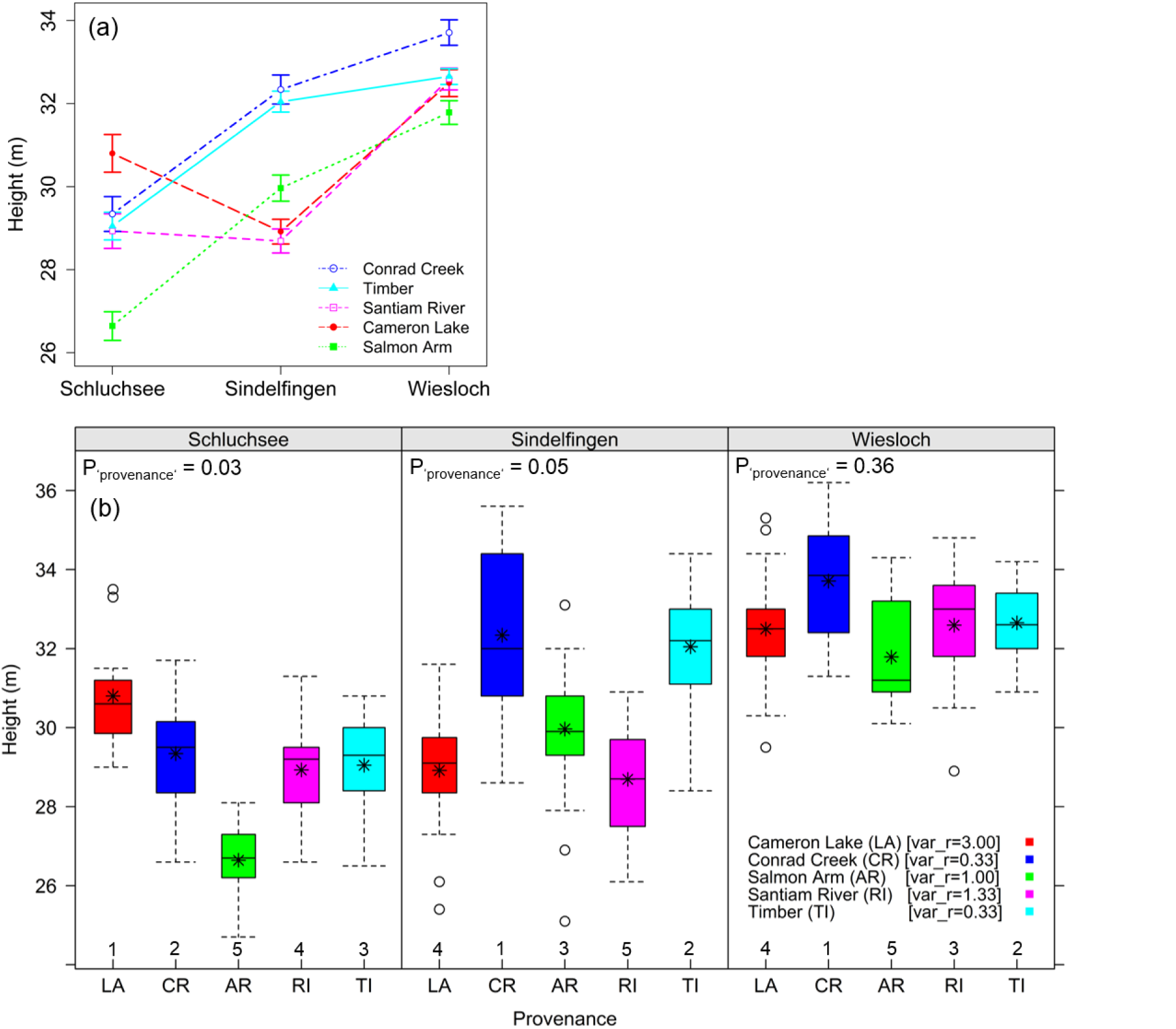
Height data of five Douglas-fir provenances growing on three sites in south-western Germany. (a) Heights (mean ± standard error, N= 9 to 34) plotted over sites to illustrate the genotype-environment-interaction effect. (b) Boxplots of heights with the mean per provenance and site depicted by an asterisk (*). The number of trees per provenance and site is indicated by the width of the boxes. The P-value for a significant provenance effect within each site is given in the top left corner of each panel. Numbers at the x-axis denote the provenance ranking within each site. As a measure of stability of provenances rankings across environments, the variance of ranks across sites (var_r) is given in the legend.

In order to test whether these differences were significant, we fitted mixed linear models to the height data, taking two different approaches. Table 5 shows the significant explanatory variables of the two models. In the provenance-based model (eq. 3), both, ‘provenance’- and ‘site’-main effects (P=0.027 and P < 0.001, respectively), as well as the interaction between these two factors (P=0.0175), were significant. To validate that these results, which are based on the set of trees analysed for both height growth and genotypes (n=315), are representative for the entire set of trees measured for height (n=598), we also fitted the provenance-based model to this larger data set. As a result, we obtained the same pattern of statistical significance, albeit with lower P-values (P-values of 0.0029, < 0.001, and 0.0013 for ‘provenance’, ‘site’, and ‘provenance’-×-‘site’ effects) demonstrating that the merged data set of 315 trees was indeed a representative subsample. The visualisation of the interaction effect and the variance of provenance ranks across sites (Figure 3) revealed that the high growth performance of ‘Conrad Creek’ and ‘Timber’ is quite stable across sites. Both provenances are among the top three ranks on all sites. By contrast, ‘Salmon Arm’ was relatively stable in its low growth performance across sites. Only on site ‘Sindelfingen’ this provenance was able to grow better (29.9 ± 0.3 m) than ‘Santiam River’ (28.7 ± 0.3 m) and ‘Cameron Lake’ (28.9 ± 0.3 m). These latter two provenances showed highly site-dependent growth, reflected in the highest variance of ranks across sites (1.33 and 3.0 respectively). While on site ‘Schluchsee’ provenance ‘Cameron Lake’ was best performing (30.8± 0.5 m), its growth was poor on site ‘Sindelfingen’ (28.9 ± 0.3 m). Similarly, provenance ‘Santiam River’ showed comparable growth on sites ‘Sindelfingen’ (28.7 ± 0.3 m) and ‘Schluchsee’ (28.7 ± 0.4 m), in contrast to the general trend of lowest growth at site ‘Schluchsee’.

**Table 5.** Significance of terms of the fixed-effects part of the models according to *F*-Tests. DFnum, degrees of freedom of the numerator; DFdenom, degrees of freedom of the denominator; Cluster-1, Cluster-2, membership to cluster 1 and cluster 2 as identified in the population genetic analysis using STRUCTURE with *K* = 5 assumed subpopulations

| Model | Model term | DFnum | DFdenom | F-value | P-value |
| --- | --- | --- | --- | --- | --- |
| Provenance based<br>(eq. 3) | Intercept | 1 | 284 | 34305.8 | < 0.0001 |
|  | Provenance | 4 | 16 | 3.64 | 0.0272 |
|  | Site | 2 | 16 | 39.0 | < 0.0001 |
|  | Site × Provenance | 8 | 16 | 3.41 | 0.0175 |
| STRUCTURE<br>based<br>(eq. 5) | Intercept | 1 | 282 | 18031.3 | < 0.0001 |
|  | Site | 2 | 282 | 19.491 | < 0.0001 |
|  | Cluster-1 | 1 | 282 | 4.574 | 0.033 |
|  | Cluster-2 | 1 | 282 | 4.91 | 0.0275 |

In order to analyse whether the differences among provenances within sites were significant, separate models per site were fitted. On sites ‘Schluchsee’ and ‘Sindelfingen’, a significant effect of ‘provenance’ was detected (P=0.03 and P=0.05, respectively), while the differences between provenances on site ‘Wiesloch’ were not significant (P=0.36).

For the two-factorial provenance-based model, the correlation between observations within the same level of ‘plot’, Corr_Plot_, was 0.37, i.e. the random effect accounted for 37%, the within-group error for 63% of the variability not explained by the fixed effects.

In the STRUCTURE-based model (eq. 5), the predictors ‘site’, membership to STRUCTURE ‘cluster-1’ and membership to STRUCTURE ‘cluster-2’ were significant. Corr_Plot_ was 0.44, i.e. the random effect accounted for 44%, the within-group error for 56% of the variance not explained by the fixed effects.

The explanatory power of the models was analysed by calculating correlations between observed and predicted values. The squared correlation between observed values and predicted values for the provenance-based model was 0.689 (Supplementary Figure 7), and 0.695 for the STRUCTURE-based model (Supplementary Figure 7).

The assumptions of independent, normally distributed residuals and normally distributed random effects were assessed graphically (Supplementary Figures 3, 4, 5). There were no indications that any of these assumptions were violated.

A significant correlation between growth and field site climatic differentiation was found based on a Mantel test (R=0.51, P=0.0006). The correlation between growth and genetic variation was significant (R=0.25, P=0.0404). In addition, the question was raised whether there was a correlation between growth and genetic differentiation after taking differentiation of field site conditions into account. To address this question, we performed a partial Mantel test. In this case, no significant correlation was found (R=0.19, P=0.0822).

### Height of Douglas-fir trees in relation to climatic conditions at the provenances origin

Moreover, significant correlations were found between genetic variation and climate of the provenance (R=0.59, P=0.0003) and between genetic variation and geographic distance between provenances (R=0.42, P=0.0009). A significant correlation between genetic variation and climatic differentiation was also found when geographic distance was taken into account by applying a partial Mantel test (R=0.46, P=0.0008). Finally, Mantel tests revealed no significant correlation either between growth and original climate of the provenances (R=0.07, P=0.2163), or between growth and geographic distance between provenances (R=-0.03, P=0.5969). Given that none of the aforementioned tests were significant, we did not perform a partial Mantel test.

## Discussion

### Genetic variation among provenances as revealed by microsatellite markers

Genetic variation at the four analysed SSR loci was high and comparable to earlier studies (Slavov et al. 2004, Krutovsky et al. 2009). A characteristic of the used markers is the presence of ‘null’ (non-amplified) alleles. These reached frequencies up to 31 % in our study. In addition to ‘null’ alleles, errors due to excessive stuttering might have occurred, thus explaining the fact that F_IS_ values remained high and significant even after genotypic adjustments. These results are comparable to the results reported by Krutovsky et al. (2009), who used the same microsatellite markers.

Different levels of genetic diversity among provenances are suggested by our results. Both the highest variability (in terms of n_a_; see Table 3) and diversity (in terms of A, H_o_ and H_e_; see Tables 3 and 4) were measured in provenances ‘Santiam River’ and ‘Timber’. The lowest variability and diversity were observed in the two northernmost provenances, ‘Salmon Arm’ and ‘Cameron Lake’, while provenance ‘Conrad Creek’ displayed median values. This pattern agrees with the glacial history and post-glacial migration of the species. Pollen and macrofossil evidence place a significant refugial population in the Willamette Valley (northern Oregon), in a region where both ‘Santiam River’ and ‘Timber’ provenances reside (Tsukada 1982, Gugger and Sugita 2010). A relatively high genetic diversity for this area, which is in concordance with the existence of a refugial population, has also been indicated by a range wide study of Li and Adams (1989). Long persistence of these populations on their current locations might have allowed for accumulation of genetic diversity. In contrast, post-glacial recolonization may have led to losses of genetic diversity due to genetic drift, resulting in a reduced genetic diversity in the northernmost provenances of our study.

A significant differentiation of ‘Salmon Arm’ from all other provenances is shown by pairwise measures of genetic differentiation, as well as by the Bayesian Analysis of population structure (based on the STRUCTURE software) and the Factorial Correspondence Analysis (FCA). Even though the number of loci genotyped in the present study is not high, we consider the presented results of the STRUCTURE analysis as robust, since Hubisz et al. (2009) reported that by using the locprior-Option, the true population structure will be well approximated with a number of loci as small as two. The STRUCTURE analysis clearly shows that irrespective of the chosen clustering solution, individuals of provenance ‘Salmon Arm’ form a distinct subpopulation. This supports the different origin of the source population, which is located in an area where the two varieties – coastal and interior – may intermingle (Kohnle et al. 2012, Gugger et al. 2010).

On the other hand, genetic differentiation among the four remaining, coastal provenances was more limited, but could also be identified by means of the aforementioned methods. In particular, provenances ‘Santiam River’ and ‘Timber’, which originate from habitats relatively close to each other in Oregon, were genetically homogenous. These two southernmost provenances could be distinguished from the northernmost coastal provenance ‘Cameron Lake’ from British Columbia. ‘Conrad Creek’, originating from a region between the two extremes, showed an admixed gene pool between the northernmost and the southernmost coastal provenances (see the clustering solution of STRUCTURE for three assumed subpopulations; *K* = 3), which is in agreement with a latitudinal gradient. Remarkably, very similar patterns of genetic differentiation were found in a complementary study, where the genetic diversity of a subset of trees of the same populations were analysed on the basis of approximately 80,000 SNP markers (Müller et al. 2015). Both isolation-by-distance (IBD) and geographic barriers acting currently and historically might account for this differentiation pattern among provenances studied in this work. Krutovsky et al. (2009) analysed the relationship between genetic differentiation and geographic distance with a larger number of populations covering the whole range of the coastal variety within the United States and found a positive and significant correlation between genetic differentiation and distance.

Besides migration and drift, an additional potential cause of the revealed genetic differentiation among the study provenances could be natural selection, leading to isolation-by-adaptation (IBA). In particular, adaptive divergence can also increase genome-wide differentiation by promoting general barriers to neutral gene flow, thereby facilitating genomic divergence via genetic drift. This latter process can yield a positive correlation between adaptive phenotypic divergence and neutral genetic differentiation (Nosil et al. 2008). Mantel tests carried out in our study showed that there was a correlation between genetic differentiation and climate at origin, even when the effects of IBD were taken into account. IBA may occur due to several ecological differences like the intensity of summer drought, or increasingly continental climate, which characterises the interior part of the region where the source populations originate from (Hermann and Lavender 1990, Nosil et al. 2008, Mosca et al. 2014). A genetic basis of drought tolerance of Douglas-fir has been recently indicated based on physiological data and growth in provenance trials (Jansen et al. 2013), as well as genetic differentiation in drought related genes (Müller et al. 2015). Additionally, a number of genes have been associated with cold hardiness in the frame of association studies (Krutovsky and Neale 2005, Eckert et al. 2009). Even if the loci of our study are considered as selectively neutral, it cannot be precluded that they carry the imprints of natural selection due to selection at closely linked genes due to genetic hitchhiking (Andolfatto 2001).

### Significant effects of ‘site’, ‘provenance’, and their interaction on height growth

The statistical analysis of height growth of Douglas-fir revealed significant differences between sites. Height growth is a well-established indicator for site productivity, that is, besides being putatively affected by the genetic background of the trees, it is largely determined by growth relevant environmental factors such as temperature, precipitation and soil properties (Puettmann et al. 2009, Messaoud and Chen 2011, Darychuk et al. 2012). Accordingly, differences in height growth among sites found in our study clearly reflect gradients in environmental factors that will determine height growth. This correlation between growth and among-site climatic variation was also confirmed by means of Mantel test and was highly significant. The site with the lowest growth potential, ‘Schluchsee’, is located at a high elevation (1050 m a.s.l.), with accordingly lower mean annual temperature (6.1 °C) and a shorter growing period, whereas the site ‘Wiesloch’ is located in the lowland (105 m a.s.l) and is characterised by a mean annual air temperature of 9.9 °C. Among the three sites studied here, ‘Sindelfingen’ is situated at a medium elevation and is characterised by intermediate growth potential as well as intermediate site conditions (8.2 °C, 490 m a.s.l.). In a study on growth phenology in seedlings of coastal Douglas-fir, Gould et al. (2012) reported increased height growth in environments with higher mean annual temperatures, regardless of seed source origin. Interestingly, the rather low annual precipitation on site ‘Wiesloch’, in combination with a sandy soil type does obviously not override the positive effect of the higher temperatures typical for this site. This is in agreement with recently published results from field trials in Austria and Bavaria (SE Germany) showing that increasing temperature at the planting site had a significant positive effect on height growth of Douglas-fir, whereas precipitation variation did not explain growth differences among sites (Chakraborty et al. 2015).

Attempts to analyse the effect of seed source origin (i.e. the effect of the factor ‘provenance’) on phenotypic traits usually assume that such differences of geographic position reflect differences of genetic variation in that trait. One objective of this study was to analyse the effect of seed source origin on height growth. Though less pronounced than the effect of field site, we showed that provenance has also an effect on growth (Table 5). This is supported by the Mantel test between phenotypic and genetic distances, too, but it is not confirmed by the partial Mantel test which takes the climatic differentiation among field sites into account. Therefore, our results do not support a link between climate of origin and growth performance which would be indicative of adaptation. However, it should be taken into account that no more than five provenances planted on three sites could be included in our study, which is probably too few to draw general conclusions. Significant effects of the environment of origin on height growth have been shown elsewhere based on larger numbers of Douglas-fir provenances (Krakowski & Stoehr 2009, Leites et al. 2012, Chakraborty et al. 2015). For example, previous reports suggested that provenances from sites with high mean annual temperature show a better growth potential (Leites et al. 2012, Chakraborty et al. 2015), which is in agreement with the constantly high growth potential of ‘Conrad Creek’ observed in our study. In contrast, provenance ‘Cameron Lake’ showed highly site-dependent growth performance. The high growth potential on site ‘Schluchsee’ along with a near-average or low height growth on sites ‘Sindelfingen’ and ‘Wiesloch’, respectively, may suggest that on the latter two sites, the low annual precipitation might restrict height growth of ‘Cameron Lake’ (climate of origin is wetter; see Table 1). Recent measurements on other sites of the provenance tests confirm these trends; ‘Conrad Creek’ was always above average, whereas the height of ‘Cameron Lake’ was site specific elsewhere, too.

### Genetic variation in microsatellite loci is related to phenotypic variation in height growth

The superior growth of either ‘Cameron Lake’ (on site ‘Schluchsee’) or ‘Conrad Creek’ (on sites ‘Sindelfingen’ and ‘Wiesloch’) may suggest a north-south gradient of growth performance for the coastal provenances, which agrees with results reported by Eilmann et al. (2013). Interestingly, the high growth performance of the provenances ‘Conrad Creek’ and ‘Cameron Lake’ coincides with a low genetic diversity of these provenances (Table 3). With the accompanying genetic analysis, we are able to show that this – pronounced and highly significant – phenotypic difference might indeed be related to a genetic differentiation (compare Figures 2, 3, Supplementary Table 5, Supplementary Figure 2). In the STRUCTURE-based modelling approach, the membership proportions to Cluster 1 and Cluster 2 were identified as significant predictors, providing statistical evidence for a genetic component in the observed phenotypic differentiation in height growth (Table 5). Cluster 1 (indicated in blue in Figure 2) largely represents the genetic differentiation of the provenance ‘Salmon Arm’ from the coastal provenances, while the membership proportions to Cluster 2 (indicated in red in Figure 2) separate Santiam River and Timber on the one hand, and Cameron Lake and Conrad Creek on the other. Given the fact that trees of provenance ‘Salmon Arm’ perform low in height growth on all three sites, it is not surprising that the membership to Cluster 1 is a significant explanatory variable for height growth. The pattern of membership proportions of the four coastal provenances to Cluster 2 indicate that genetic differentiation correlates with the significant differences in height growth observed between ‘Conrad Creek’ and ‘Santiam River’.

An interesting result is the quasi-identical explanatory power of the two modelling approaches. While the provenance-based model can only account for among-provenance variation, the STRUCTURE-based model potentially reflects both among-and within-provenance genetic variation. However, the quasi-identical explanatory power of the two models argues for a low extent of within-provenance genetic variation in the analysed loci. Given the putative nature of height growth as a quantitative trait, the relatively small number of microsatellite loci genotyped in this study, the large genome size of Douglas-fir (O’Brien et al. 1996) and the rapid decay of linkage disequilibrium in forest trees (Neale and Kremer, 2011) it appears reasonable that genetic variation underlying the observed differences in height growth is only partially captured by the genetic parameters derived from the SSR genotypes. As a consequence, the STRUCTURE-based model does not outperform the provenance-based approach. With the set of single nucleotide polymorphism markers developed for Douglas-fir recently (Müller et al. 2012; Müller et al. 2015), we now have the tools available to analyse the effect of adaptive genetic variation on height growth (and other phenotypic traits) directly.

## Conclusion

Considering the importance of genetic diversity for the adaptability of tree populations, and the fact that our results on height growth suggest a tradeoff between genetic diversity and growth, this study is of significance for forest managers. For a sustainable management of forest ecosystems under conditions of a changing climate, it is of particular significance to learn more about adaptedness and adaptability of tree species on sites where they are introduced. In our study, we did this by assessing phenotypic variation along environmental gradients and by relating it to climate of origin and genotypic variation. An innovative aspect of the study was to directly test a genetic component in height growth variation by deriving explanatory variables from molecular data. The use of growth data from stands of an age of over 50 years – higher than most provenance research studies known to us – increases the significance and confidence of the results. On the other hand, this study was restricted to five provenances and three sites. In order to enable the development of climate-and genotype-sensitive models of height growth for Douglas-fir, future research should be based on larger sets of sites, provenances, and genetic markers.

## Acknowledgements

The study sites are part of the International Douglas-fir provenance trial, initiated by Prof. R. Schober after a decision of the Unit of Forest Growth of the German Association of Forest Research Stations (DFFVA). The German federal state of Baden-Württemberg joined this provenance trial series in 1961 with test plantations at 15 locations, of which ten are currently still active and supervised by the Forest Research Institute of Baden-Württemberg (FVA). Seeds for the experiments were collected in 1955 by B. Strehlke from autochthonous stands in North America and from selected cone-bearing stands growing in Southwestern Germany judged by forest practitioners to display vigorous growth. We thank Dr. Axel Albrecht for helpful discussions. This study was financially supported by the Forest Research Institute Baden-Württemberg with funding to I.E., U.K., H.W. and by the German Science Foundation (DFG) with grants to U.K. (KO 931/8-1) and I.E. (EN 829/4-1) as part of the collaborative project DougAdapt (PAK 583/468).

